# Identification of Potential Oral Cancer-Specific Biomarker in Oral Potentially Malignant Disorders for Early Detection of Malignancy: A Systematic Review

**DOI:** 10.1101/2025.05.18.651790

**Authors:** Dhananjay Dhar Dwivedi, Pragya Tiwari, Yasmin Bano, Pramod Kumar, Chander Prakash Yadav, Dinesh Kumar Singhal

## Abstract

Oral malignancies are increasing and significantly contribute to cancer-related mortality. Oral Potentially Malignant Diseases (OPMDs) share several characteristics with Oral Cancer (OC) and have the potential to undergo malignant transformation. Beyond clinical and visual similarities, OC and OPMDs also exhibit common molecular alterations at the genomic and transcriptional levels. Identifying commonly mutated genes between OC and OPMDs may enhance early detection strategies. Therefore, we conducted a systematic review analyzing mutated genes from 25 studies (selected from 331 studies using the PRISMA flowchart) that investigated gene mutations in OC and OPMDs through whole-exome sequencing analysis. Across these studies, we identified a total of 7,564 mutated genes: 6,885 from OC studies and 579 from OPMD studies. Of these, 311 genes were found to be common between OCs and OPMDs. To further assess their relevance, these genes were categorized into High (N=5), Medium (N=20), and Low-frequency (N=386) groups, and their distribution was compared between OC and OPMD studies. Based on these comparisons, four genes: TP53, PIK3CA, FAT1, and TTN were identified as showing comparable mutation patterns between OC and OPMDs, suggesting their potential role as early biomarkers for malignant transformation. Furthermore, FAT1 and TTN may serve as promising candidates for assessing the likelihood of Oral Cancer development in OPMDs.

## Introduction

Oral Potentially Malignant Diseases (OPMDs) refer to “any oral-mucosal abnormality that is associated with an increased risk of developing oral cancer” (1). The OPMDs include oral leukoplakia (OLK), oral erythroplakia, oral submucous fibrosis (OSMF), oral lichen planus (OLP), etc. OPMDs share several characteristics with Oral Squamous Cell Carcinoma (OSCC) and have the potential to undergo malignant transformation. It can be associated with oral cancer-related mortality due to a lack of specific screening programs or delayed diagnosis.

More than 90% of oral cancers are OSCCs that have OPMDs-like symptoms (2,3). Worldwide, OCs have been a frequent malignancy in low- and middle-income countries with bad prognosis, aggressive behavior, and the propensity of lymph node metastasis (4). Inadequate knowledge of OC’s risk factors and low screening rate of OPMDs contribute to a higher rate of advanced-state oral cancer detections (5,6) and most of the OC cases advanced to stage III or IV due to late diagnosis (7). OSCCs have a significant impact on health and the global economy (8). The 5-year survival rate improves from 40% to 80%, if OC is detected at early stages (I and II) rather than advanced stages (9). Moreover, OPMDs have the potential to transform into OCs, which must be assessed for early malignancy detection.

Various methods are being used for OCs or OPMDs diagnosis, visual oral examination is the primary screening method which depends on the expertise and experience of the physician, and several symptoms like red and white lesions are indistinguishable (10). Various shortcomings of VOE results which depend on expertise, calibration of technique, methodology, and indistinguishable symptoms of OCs and OPMDs have been highlighted in the Global Oral Cancer forum-2016 (11). Molecular biology techniques have revolutionized diagnostics and made it possible to detect malignancies and other diseases and the advent of genome sequencing has been proven to provide substantial breakthroughs for mutations and other genomic information identification (12). Moreover, mutations are attributable to cancer whether it is inherited or acquired, and can be detected using a mutational landscape. The presence of CDKN2A germline mutations has been attributed to a higher risk of OCs (13). Tp53, CDKN2A, pRB, NOTCH1-4, EGFR, and RAS mutated genes have been detected in OCs. The altered genetics loci 3q26.32 (*PIK3CA*) 9p21.3 (*CDKN2A*), 9q34.3 (*NOTCH1*), and 17p13.1 (*TP53*) have been linked with chewing tobacco OCs (14). Various mutated genes have been reported in OCs and OPMDs which can provide a way towards early diagnosis of malignancies. The current systematic review has been performed to identify the most comparable mutation patterns between OC and OPMDs for suggesting their potential role as early biomarkers for malignant transformation.

## Materials and Methods

### Prospero Registration

The study is registered in “Prospero (International Prospective Register of Systematic Reviews) under the ID number CRD42024580222 and has been conducted as per the PRISMA guidelines.

### Literature Search

Studies that performed whole exome sequencing to identify mutated genes in Oral Cancer (OC) and Oral Potentially Malignant Disorders (OPMD) patients were searched across three databases: PubMed, Embase, and Web of Science. The study selection followed PRISMA guidelines, adhering to predefined inclusion and exclusion criteria.

### Search Terms

The following search terms were used to identify relevant studies across the selected databases: PubMed, Embase, and Web of Science.

**#1** (“cancer, mouth” OR “intraoral cancer” OR “mouth mucosa cancer” OR “oral cancer” OR “oral cavity cancer” OR “mouth cancer” OR “oral lesion, precancerous” OR “oral lesion, premalignant” OR “oral potentially malignant disease” OR “oral pre-malignant lesion” OR “oral precancerous condition” OR “oral precancerous lesion” OR “oral premalignant lesion” OR “potentially malignant oral disease” OR “potentially malignant oral disorder” OR “precancerous oral lesion” OR “precarcinoma, mouth” OR “premalignant oral lesion” OR “oral potentially malignant disorder”)

**#2** (“exome sequence analysis” OR “exome sequencing” OR “exomic sequencing” OR “WES (whole exome sequencing)” OR “WES analysis” OR “whole exome sequencing”)

### #1 AND #2 (date)

The above-given search terms and strategy were validated by the subject expert and independently reviewed and repeated by the reviewer of the studies to ensure appropriateness and completeness.

### Inclusion Criteria

1. Studies reported whole exome-sequencing-based mutations in human oral cancer and OPMDs-related studies with controls
2. Studies must be in English
3. Studies without any intervention or treatment to the cancer patients

### Exclusion Criteria

1. Studies not having control samples
2. Studies have not reported mutated gene names
3. Animal-based studies have also been excluded

### Selection Process

Two independent reviewers (DDD and PT) reviewed studies from different platforms and third and fourth reviewers (DKS and CPY) resolved any discrepancies and disagreements. The first level of screening was performed based on the title and abstract. Subsequently, the screened articles underwent full-text review, and valid reasons were recorded for each excluded article.

### Data Extraction

Data from selected studies were extracted using a data extracting form and entered in MS Excel. Along with the study title, year, geography, and the names of all mutated genes were recorded from each selected study.

### Classification and categorization of extracted genes

All extracted genes from the selected studies were classified into three categories: high, medium, and low-frequency genes, based on the number of studies in which they appeared. If a gene appeared multiple times within the same study, it was counted only once. Genes present in approximately 50% or more of the studies were classified as high-frequency, those appearing in 20% to 50% of the studies were categorized as medium-frequency, and genes found in less than 20% of the studies were labelled as low-frequency. Additionally, the percentage occurrence of each gene was compared between OC and OPMD studies. Based on this comparison, genes were further categorized as either comparable or non-comparable, depending on whether their frequency distribution was similar across both groups (OC and OPMD).

### Identification of potential biomarkers for OPMD

High and medium-frequency genes that showed comparable occurrence between OC and OPMD groups, can be considered for *risk factor stratification* for OMPD. Furthermore, these genes were analyzed using the cBioPortal for Cancer Genomics, utilizing data from TCGA (https://www.cbioportal.org/). The frequency of mutations, as well as gene alterations such as loss or gain of function, were assessed to understand their potential role in tumorigenesis.

### Data Analysis

All data analysis and plot development were performed using R 4.4.1 software.

## Results

A literature search was conducted using three databases: PubMed, Embase and Web of Science, yielding 331 articles. After removing duplicates, 233 articles were screened for title and abstract and 42 studies were included for full text review. Of these 42 studies, 25 studies were finally included in the review, 19 studies focused on Oral Cancer (OC) and 6 studies were on potentially malignant disorders (OPMDs) while 17 articles were excluded based on serval reasons: Assessing the effect of tobacco project (3), based on mice model (1), did not meet the inclusion criteria (7), mutation information was not available (3), TCGA based study (2), and targeted exome sequencing was used (1) (Figure 1).

**Figure 1:**
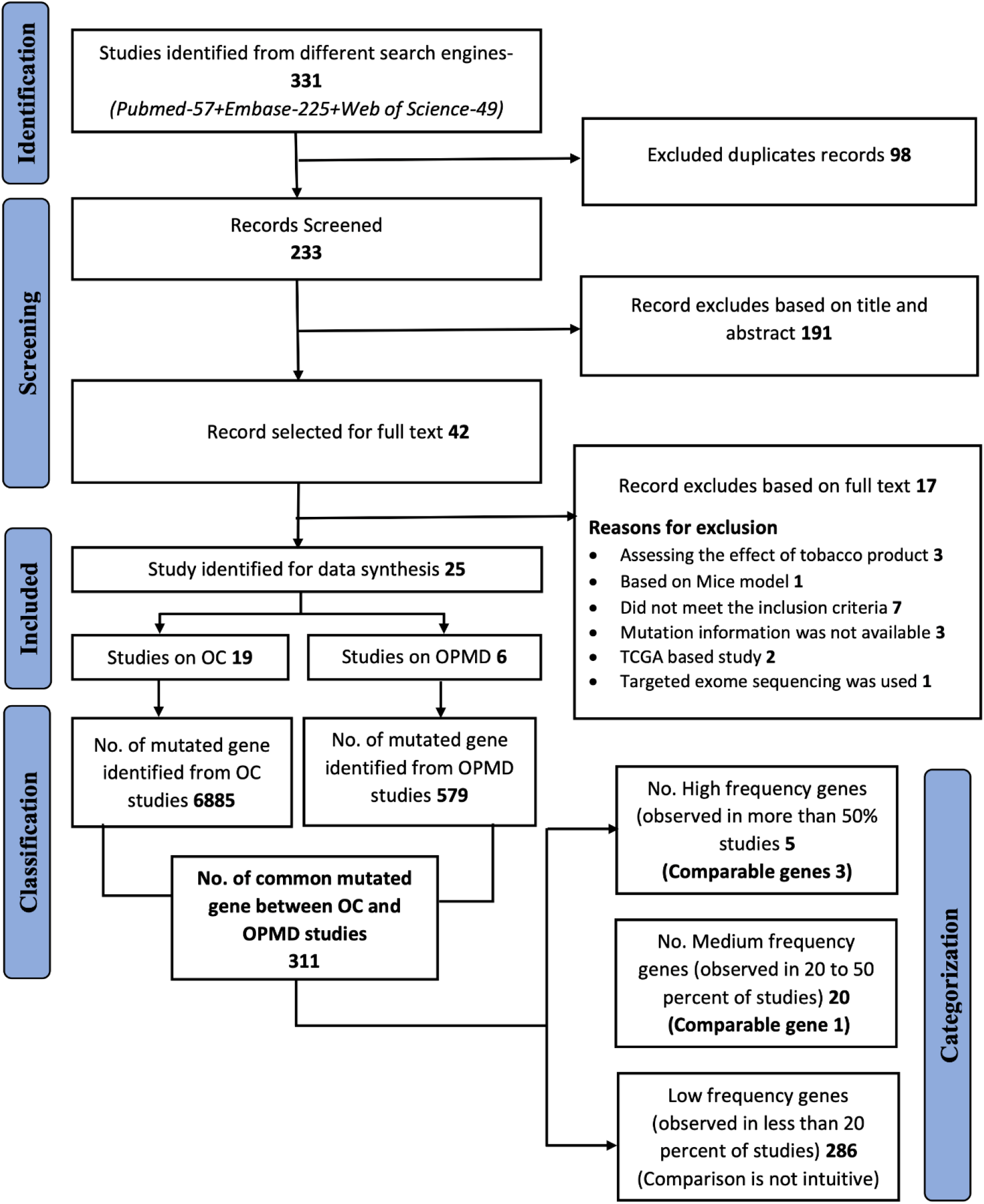
Study flow chart in accordance with PRISMA.

A total of 7,464 mutated genes were reported across the studies, with 6,885 genes identified in oral cancer patients (from 19 studies) and 579 genes in oral potentially malignant disorder (OPMD) patients (from 6 studies). Among these, 311 mutated genes were common between OC and OPMD studies. These genes were categorized into three frequency-based groups: high-frequency genes (N=5), medium-frequency genes (N=20), and low-frequency genes (N=286). A gene was classified as high frequency if it was present in ∼=50% of the studies, regardless of whether it appeared in OC or OPMD studies. Genes observed in 20 to <50% of studies were classified as medium-frequency, while those found in less than 20% of studies were categorized as low-frequency genes (Figure 1).

After the initial classification, each gene within the high, medium, and low-frequency groups was further categorized as either comparable or non-comparable by comparing its distribution in OC and OPMD studies. This classification was based on the difference in the gene’s occurrence percentage between the two groups. For example, the gene NOTCH1 was identified in 15 out of 25 studies, meaning it was present in 60% of the studies, classifying it as a high-frequency gene. When examining its distribution across OC and OPMD studies, NOTCH1 was found in 13 out of 19 OC studies (68.4%) and only in 2 out of 6 OPMD studies (33.3%). This significant difference in prevalence between the two groups suggests that NOTCH1 exhibits a notable disparity in its occurrence between OC and OPMD cases, categorizing it as a non-comparable gene. Likewise, all the other genes were also defined on a similar pattern.

The Five genes: NOTCH1 (60%, 15 out of 25 studies), TP53 (60%, 15 out of 25), PIK3CA (56%, 14 out of 25), FAT1 (52%, 13 out of 25), CDKN2A (48%, 12 out of 25) and were identified as high-frequency genes, appearing in more than ∼50% of the studies. Among these, the proportions of FAT1 (52.6% in OC vs. 50% in OPMD), PIK3CA (58% in OC vs. 50% in OPMD), and TP53 (63% in OC vs. 50% in OPMD) were comparable between OC and OPMD studies. In contrast, NOTCH1 (68% in OC vs. 33% in OPMD) and CDKN2A (53% in OC vs. 33% in OPMD) were more prevalent in OC studies (Table 1).

**Table 1:** Comparison of high-frequency genes (observed in more than 50% of studies) in Oral Cancer and OPMD studies.

| Gene Name | Oral Cancer |  |  | OPMD |  |  | Overall |  |  |
| --- | --- | --- | --- | --- | --- | --- | --- | --- | --- |
|  | No. of studies included | No. of studies in which gene observed | % | No. of studies included | No. of studies in which gene observed | % | No. of studies included | No. of studies in which gene observed | % |
| NOTCH1 | 19 | 13 | 68.4 | 6 | 2 | 33.3 | 25 | 15 | 60.0 |
| TP53 | 19 | 12 | **63.2** | 6 | 3 | **50.0** | 25 | 15 | 60.0 |
| PIK3CA | 19 | 11 | **57.9** | 6 | 3 | **50.0** | 25 | 14 | 56.0 |
| FAT1 | 19 | 10 | **52.6** | 6 | 3 | **50.0** | 25 | 13 | 52.0 |
| CDKN2A | 19 | 10 | 52.6 | 6 | 2 | 33.3 | 25 | 12 | 48.0 |

Twenty genes: CASP8, TTN, LRP1B, HRAS, DNAH5, PLEC, ZFHX4, COL22A1, MUC4, ARID2, ATP10A, DNAH9, FAT2, HLA-B, KRAS, MAMLD1, PCLO, PTCHD2, PTEN, and RECQL4 were classified as medium-frequency genes. Among them, only one gene i.e. TTN, exhibited similar proportions in both OC and OPMD studies, indicating comparable prevalence between the two groups (Table 2).

**Table 2:**
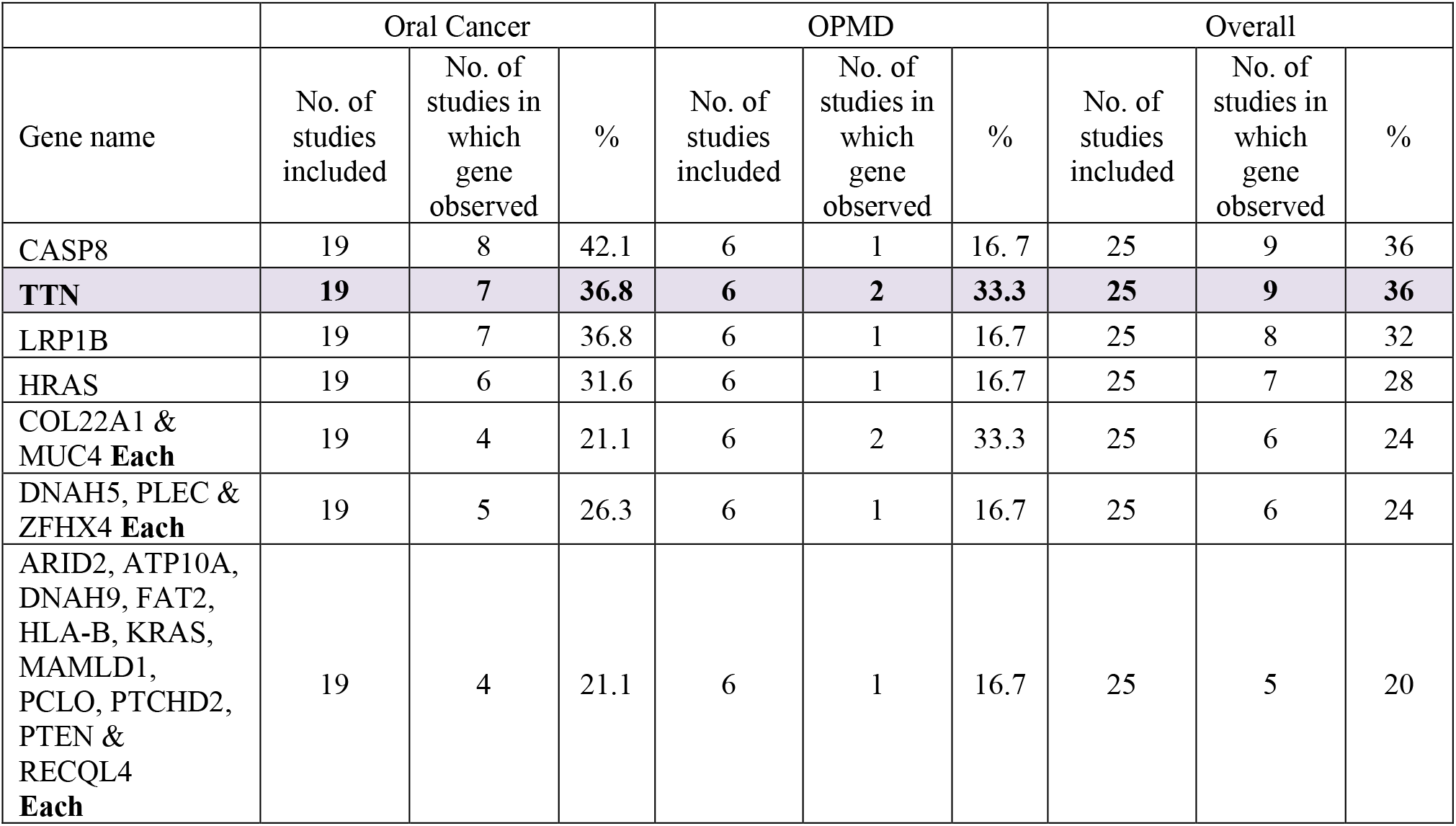
Comparison of Medium frequency genes (observed in ≥ 20% and <50% of studies) in Oral Cancer and OPMD studies.

|  | Oral Cancer |  |  | OPMD |  |  | Overall |  |  |
| --- | --- | --- | --- | --- | --- | --- | --- | --- | --- |
| Gene name | No. of studies included | No. of studies in which gene observed | % | No. of studies included | No. of studies in which gene observed | % | No. of studies included | No. of studies in which gene observed | % |
| CASP8 | 19 | 8 | 42.1 | 6 | 1 | 16.7 | 25 | 9 | 36 |
| **TTN** | **19** | **7** | **36.8** | **6** | **2** | **33.3** | **25** | **9** | **36** |
| LRP1B | 19 | 7 | 36.8 | 6 | 1 | 16.7 | 25 | 8 | 32 |
| HRAS | 19 | 6 | 31.6 | 6 | 1 | 16.7 | 25 | 7 | 28 |
| COL22A1 & MUC4 **Each** | 19 | 4 | 21.1 | 6 | 2 | 33.3 | 25 | 6 | 24 |
| DNAH5, PLEC & ZFHX4 **Each** | 19 | 5 | 26.3 | 6 | 1 | 16.7 | 25 | 6 | 24 |
| ARID2, ATP10A, DNAH9, FAT2, HLA-B, KRAS, MAMLD1, PCLO, PTCHD2, PTEN & RECQL4 **Each** | 19 | 4 | 21.1 | 6 | 1 | 16.7 | 25 | 5 | 20 |

Similar to high- and medium-frequency genes, 286 genes fall under the low-frequency gene category. Due to their low occurrence, a meaningful comparison between OC and OPMD is not feasible. These low-frequency genes are summarized in (Supplementary Table S1).

In a nutshell, there are four genes: TP53, PIK3CA, FAT1, and TTN were identified as showing comparable mutation patterns between OC and OPMDs, suggesting their potential candidature as early biomarkers for malignant transformation. These genes were further analyzed using the cBioPortal for Cancer Genomics, utilizing data from TCGA. The frequency of mutations, as well as gene alterations such as loss or gain of function, were assessed to understand their potential role in tumorigenesis. The mutations in TP53, PIK3CA, FAT1, and TTN genes affect various HNSC cancer cases (Table 3). The mutations in TP53 and TTN affect more than 70% and 44 % of HNSC whereas PIK3CA affects more than 68% of cases by copy number variants associated gain or loss. FAT1 affects more than 22% of cases by mutations and more than 35% by CNV-associated gain of function (Table 3).

**Table 3.**
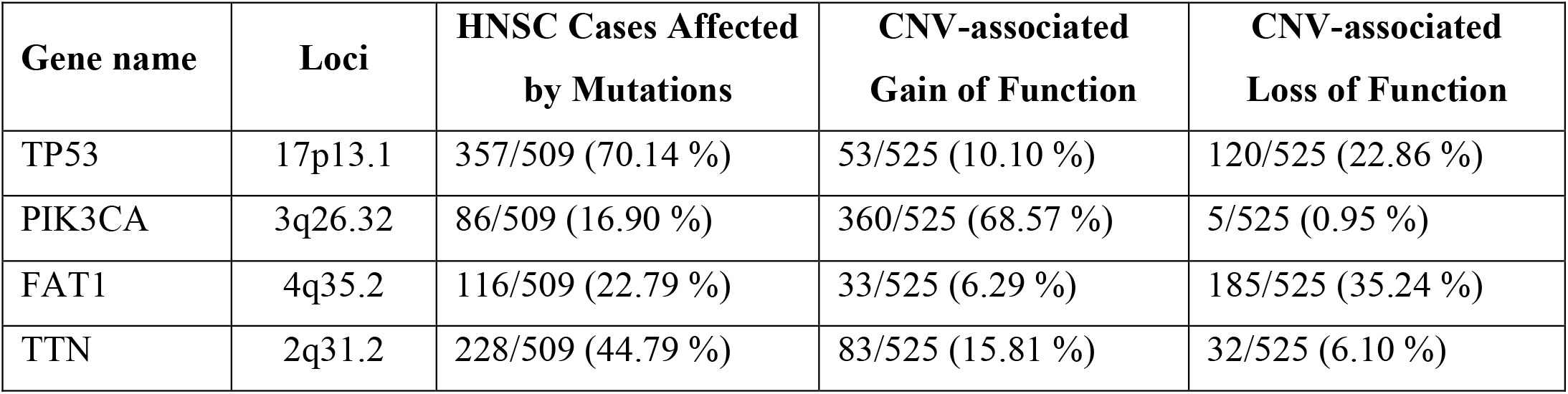
Mutations, gene alterations in four comparable genes between OCs and OPMDs from TCGA data.

## Discussion

OPMDs comprise a group of oral conditions with a risk of transformation into OCs. Their clinical presentation often overlaps with OCs in terms of morphological parameters, highlighting the need to explore molecular markers for improved disease diagnosis. In this study, we investigated commonly mutated genes in OC and OPMD patients to assess their potential role as early biomarkers for malignant transformation. A total of 311 genes were found to be shared between OC and OPMD patients. Among these, five genes: NOTCH1, TP53, PIK3CA, FAT1, and CDKN2A were identified in approximately 50% of the studies. When comparing their prevalence between OC and OPMD, three genes: TP53, PIK3CA, and FAT1 exhibited similar distributions between the two groups (Figure 2), whereas NOTCH1 and CDKN2A were more frequently detected in cancer patients. Additionally, 20 genes were identified in ∼20–30% of the studies, with TTN displaying a comparable distribution between OC and OPMD patients. Furthermore, eleven genes: ARID2, ATP10A, DNAH9, FAT2, HLA-B, KRAS, MAMLD1, PCLO, PTCHD2, PTEN, and RECQL4 also showed similar mutation patterns between OC and OPMD. However, due to the limited number of studies, no definitive conclusions can be drawn regarding their significance. The remaining 286 genes were very infrequent to compare their distribution between OCs and OPMDs.

**Figure 2:**
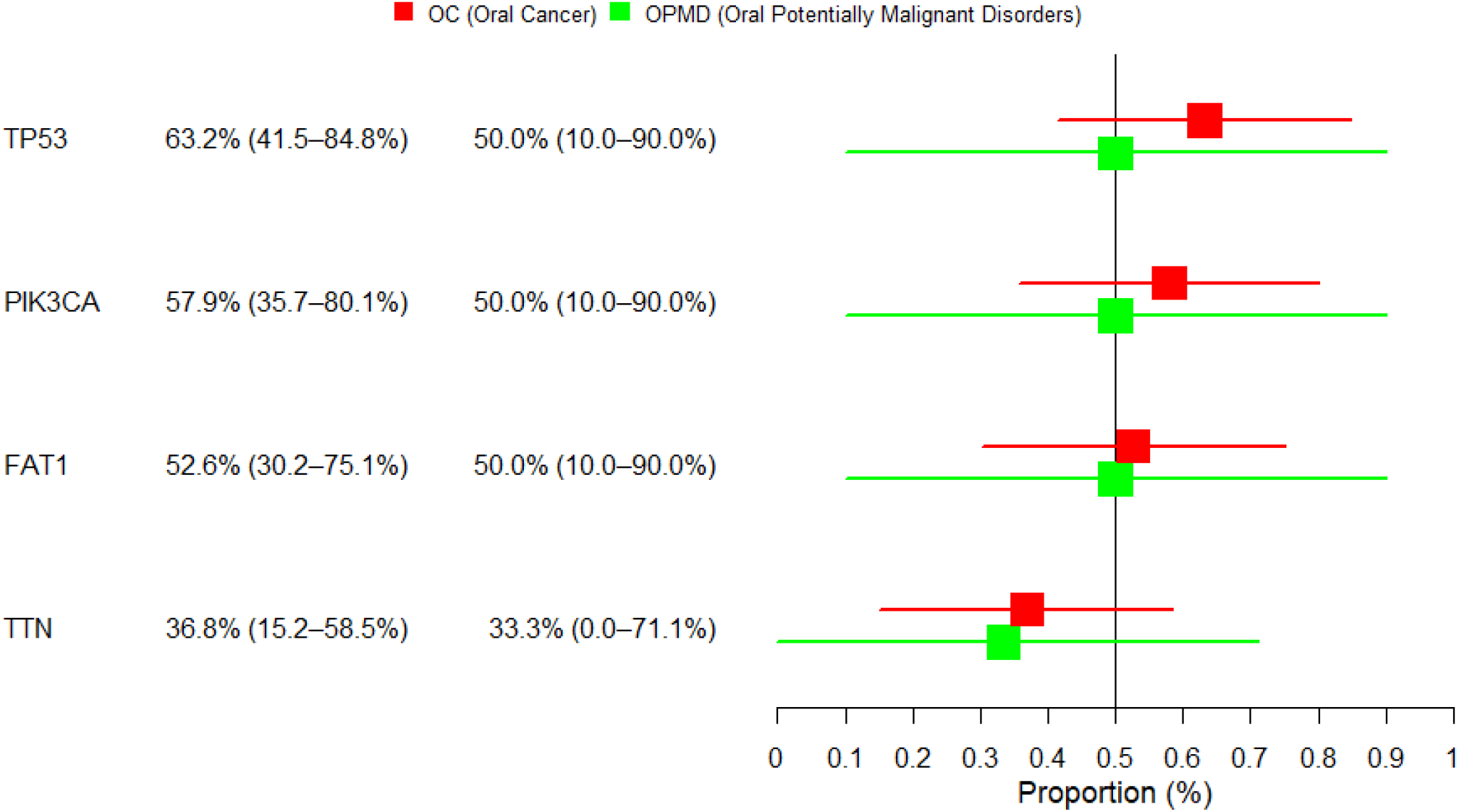
Distribution of Potential Oral Cancer-Specific Gene in Oral Potentially Malignant Disorders.

As per the literature, FAT1 and TTN may serve as promising candidates for assessing the likelihood of Oral Cancer development in OPMDs. The regulation of TP53, NOTCH1, and CDKN2A genes along with other factors play a role in cancer control or progression (15,16). TCGA-based analysis revealed that 864 NOTCH1 mutations affected 812 cases (18.37% Head and Neck cancer cases) among 41 projects (Table 3 & Supplementary File 1), and 4051 CNV events in 4621 cases affected loss and gain of function for NOTCH1. [NO_PRINTED_FORM] identified mutations in 409 cancer-related genes from the oral tissue and normal tissue of 47 Japanese patients, in which *TP53* (61.7%), *NOTCH1* (25.5%), *CDKN2A* (19.1%), and *PIK3CA* (10.6%), genes were found to be mutated (17). NOTCH1 mutations were identified in 54% of OSCC and 60% of leukoplakia Chinese patients that overlapped with the mutations reported in Caucasians and found altered regulatory domain attributed to gain of function (18). Mutated NOTCH1 (31%) and TP53 (77%) 13 oral cancer patients (19). NOTCH1 mutations have been found in 44 out of 168 paraffin-fixed OSCC samples, which are associated with its elevated expression (20).

TP53 has been studied widely as a malignancy marker and is considered a prominent marker for cancer. TP53-SNPs have been attributed to OSCC and OPMDs in a PCR-based study in Argentine patients (21). The mutated TP53 in OLP has been considered a risk factor for the transformation and progression of the disease (22).

TCGA data showed 16.90% of HNCC cases affected by the PIK3CA mutated gene and CNVs contribute to the gain and loss of functions (Table 2). The 4% mutations in the PIK3CA gene are associated with OSCC development in the Indian population (23). PIK3CA codes for PI3K and its mutations are reported to activate PI3K-AKT-mTOR signalling that causes sustained cell growth and cell invasion in HNSC (24). The presence of PIK3CA mutations in OSCC and salivary DNA suggested its implication for non-invasive ways of cancer prognostics and follow-up care (25).

FAT1 SNP-variant has been found in OPMDs and ORCs in males with or without substance use (26). The TCGA-based data analysis revealed loss and gain of function of FAT1 and 22% of cases were affected due to the presence its mutations. FAT1’s loss of functions has been associated with epithelial-mesenchymal transition modulation and stemness in cancer cells and its expression was reported upregulated in OCs (27,28). In oral leukoplakia, patients genomic landscape revealed the presence of mutations in FAT1 (20%), NOTCH1 (13%), and TP53 (28%) OCs-associated genes (29). Mutated FAT1 and LRP1B genes have been found in high tumor mutational burden patients and are also associated with chemoresistance in head and neck cancer patients (30).

CDKN2A, a kinase that regulates cellular growth and division, was found mutated in various cancers and specifically found mutated in more than 20% of HNSCC cases at TCGA, and its CNVs cases gain function in 55.01% of cases. CDKN2A/P16^INK4a^ altered expression in OPMDs has been correlated with a two-fold increase in the rate of transformations (31). SNPs were found in 20% OPMDs and 30% OCs cases in CDKN2A’s 1^st^, 2^nd^, 3^rd^-exons-specific sequencing (32). The germline mutations in CDKN2A are evident for the development of HNSCC and impute risk factor consideration for the disease (33). Targeted whole-exon sequencing of TP53, HRAS, PIK3CA, NOTCH1, CDKN2A, FBXW7, and BRAF genes of 41 cases including OLK, OLP revealed the presence of oral cancer driver gene mutations in four OLK and one OLP cases, which were located in tongue, attributed to a higher risk of transformation and as compared to the other sites (34).

CASP8 modulates apoptosis and has been reported mutated in various types of cancer but here role is not understood yet. [NO_PRINTED_FORM] reported 56 % OCs and 30% leukoplakia cases CASP8 mutation-positive (35). The OCs showed more mutated allelic frequency as compared to leukoplakia cases and loss of CASP8 activity enhances the rate of oral tumor 4-nitroquinoline-1-oxide, a carcinogen in Casp-/-mice. Casp8 mutations and CNV-associated loss or gain of functions affect the variably in HNSC cases (Supplementary Table S2.).

In TCGA-based analysis mutated TTN affected more than 44% of HNSCC cancers, and *TTN* SNP rs10497520 has been considered a potential genetic marker in OSCC (36). TTN/TP53 co-mutation has been attributed to the chemotherapy response in lung cancer patients (37). Mutation in TTN affects angiogenesis in ovarian cancer and can be used as a genetic marker for ovarian cancer detection (38).

Furthermore, LRP1B has been considered as a tumor suppressor gene and found to be altered due to genetic and epigenetic changes in various cancers (39). LRP1B polymorphism has been demonstrated as a prognostic marker in OSCC with diabetes mellitus (40). LRP1B has been predicted as an HPV integration site with a frequency of 5.8% in cervical cancer and it has been considered a prognostic marker and attributed to poor disease outcomes in HNSCC (41,42). In TCGA-based analysis, more than 9% of HNSC cases showed loss of LRP1B expression, which has been directly correlated with enhanced chemo and radioresistance in HPV-positive cases (43). Very few studies have been carried out on the longest-known proteins and need to be explored.

## Conclusion

Our systematic review highlights the shared molecular landscape between Oral Cancer (OCs) and Oral Potentially Malignant Diseases (OPMDs), identifying 311 commonly mutated genes. Among these, TP53, PIK3CA, FAT1, and TTN exhibit comparable mutation patterns in both OC and OPMDs, underscoring their potential as early biomarkers for malignant transformation. FAT1 and TTN emerge as promising candidates for assessing the probability of OPMDs progressing to OC. These findings emphasize the importance of genetic screening in early detection strategies and provide a foundation for future research to validate these biomarkers in clinical settings.

## Supporting information

https://acrobat.adobe.com/id/urn:aaid:sc:AP:9f56ff03-4a9c-4387-a703-ded2941177e5

https://acrobat.adobe.com/id/urn:aaid:sc:AP:073fb316-ae45-4717-8744-1c3f3d48940c

## Author Contributions

**DDD:** Data curation; Formal analysis; software. **PT:** Data curation; Formal analysis; software. **YB:** Formal analysis; writing – review and editing; Validation. **PK:** Conceptualization; formal analysis; Validation. **CPY:** Conceptualization; writing – review and editing; software; Validation; supervision. **DKS:** Conceptualization; writing – review and editing; software; Validation; supervision.

