## Supplementary material for "Identification of Potential Oral Cancer-Specific Biomarker in Oral Potentially Malignant Disorders for Early Detection of Malignancy: A Systematic Review": https://acrobat.adobe.com/id/urn:aaid:sc:AP:9f56ff03-4a9c-4387-a703-ded2941177e5

**Supplementary Table S1: Comparison of low-frequency genes (observed in less than 20% of studies) in Oral Cancer and OPMD studies**

| Gene name | OC (N=19) | OPMD (N=6) |
|---|---|---|
| APC, ATM, ATP7B, CELSR2, NGA4, DNAH1, NAH12, DNAH3, DNAH6, EGFR, EGR3, EP300, EPHA2, FAM120B, FAM186A, FRMPD1, HDAC6, JAK2, KCNMA1, KMT2B, LRP2, MYBBP1A, NOTCH3, NUMA1, NUP210L, PCDH10, PRRC2B, RANBP2, ROBO2, SUPT6H, TAOK2, TNRC18, TSC2, USP9X, WNK2, XDH, ZNF469, ZNF717, ZNF831, & ZZEF1 **each** | 3 (15.8) | 1 (16.7) |
| ACAD11, AHNAK2, ANK1, ANKRD36, ATP2A1, ATP8B3, CADPS, CATSPERG, CDC25A, CDC42, COL12A1, COL5A3, CTNNB1, DCAF12L2, DKC1, DNAH14, DNAJC13, EIF2AK3, ERBB4, ERN2, FAM83C, FBXW7,FGFR3, FHAD1, FOXJ2, GLI2, GNAS, GZF1, HIST1H1E, HSPG2, IDH2, IPO8, KIAA1549L, KIF7, KIT, LENG8, LRP5, MAP4K1, MERTK, NTRK3, OR5I1, PCDHGA2,PDILT, PKD1,PLD2, PLIN4, PLXNB2, POLR2A, POTED, POTEF, PRPF4B,RAG2, RBM12B, SBNO2, CN2A, SKOR2, SLC34A2, SMG7, STK11, SUDS3, SYT16, TET2,TIMD4, TRIM10, UNC13C, VWA5B1, ZNF236, & ZNF318 **each** | 2 (10.5) | 1 (16.7) |
| BAIAP3, BEST4, CDH6, PLIN3, SHC3, & SPAG17 **each** | 1 (5.3) | 2 (33.3) |
| ABCC1, ABCD2, ABL1, ADAMTS13, ADAMTS3, ADAMTSL3, AIRE, ALDH1L1, ALK, NKH,ANKLE1, ANXA11, AP2A1, AP2B1, APEX2, APOL3, ARFIP1, ARHGEF28, ARPP21, ARSI, B3GAT1, BAI3, BPTF, BRCA1,BST2, BTN3A3, C14orf37, APN3, CCDC150, CD93, CDH7,CELSR1, CENPE, CLIP2, CLUH, CNN2, COL11A2, COL16A1, COL5A1, CSNK2A2, CYFIP1, DAB2, DAB2IP, DAG1, DAZAP1, DCP1A, DCPS, DNAH7, DSEL, DYNC1H1 , DYNC2H1, EFHC1, EPG5, FAM13C, FASTKD5, FBLN1, FGFR2, FLT3, FOXP1, GAS8, GMIP, GNA11, GNAQ, GRIN2A, GRM2, HNRNPUL1, HPN, HTRA3, IL16, IL17RC, IPO5, ITGA11, ITGAL, KDR, KIAA0319L, KIAA0368, KIAA0895, KIF23, KIR2DL3, KRT4, KRT73, KRTAP10-6, LAS1L, MAP1S, MAP2, MAP3K4, MBD1, MCTP2, MET, MGAT5, MIB1, MIER1, MLH1, MON2,MPND, | 1 (5.3) | 1 (16.7) |

| |
|---|
| MTURN, MUC6, MYH6, N4BP3, NKX2-6, NLRP13, PAPOLG, PATL2, PCDHA10, PCDHA8, PCDHA9, PCDHB13, PEAR1, PEX13, PGR, PHLDB1, PIK3R3, PITPNM2, PNPLA5, PODNL1, POTEI, PRG4, PTPN11, PTPRZ1, RALGAPA1, RANGAP1, RB1, RBMXL2, RCE1, REPS1, RET, RIF1, RIMS1, RP1, RP9, SAR1B, CP2D1, SEMA4B, SEMA4F, SF3A1, SKI, SLC35E4, SLC6A17, SLC8A3, SLC9A7, SLFN11, SMAD4, STIP1, SYNPO2L, SYTL2, TAS2R31, TFRC, TLR9, TMC3, TOMM34, TOP3A, TRPM3, TTC17, TTC40, TUBGCP2, UGT2B10, USP36, VAV1, VHL, VPS13C, VSTM2B, WISP1, WNT4, ZAN, ZC3H12B, ZCCHC5, ZMIZ2, ZNF142, ZNF292, ZNF449, ZNF668, & ZNF835 **each** |

**Supplementary Table S2: Mutations, gene alterations in some comparable genes between OCs and OPMDs from TCGA data**

| Gene name | Loci | HNSC Cases Affected by Mutations | CNV-associated Gain of Function | CNV-associated Loss of Function |
|---|---|---|---|---|
| NOTCH1 | 9q34.3 | 94/509 (18.47 %) | 148/525 (28.19 %) | 68/525 (12.95 %) |
| TP53 | 17p13.1 | 357/509 (70.14 %) | 53/525 (10.10 %) | 120/525 (22.86 %) |
| PIK3CA | 3q26.32 | 86/509 (16.90 %) | 360/525 (68.57 %) | 5/525 (0.95 %) |
| FAT1 | 4q35.2 | 116/509 (22.79 %) | 33/525 (6.29 %) | 185/525 (35.24 %) |
| CDKN2A | 9p21.3 | 103/509 (20.24 %) | 64/525 (12.19 %) | 289/525 (55.05 %) |
| CASP8 | 2q33.1 | 53/509 (10.41 %) | 50/525 (9.52 %) | 79/525 (15.05 %) |
| TTN | 2q31.2 | 228/509 (44.79 %) | 83/525 (15.81 %) | 32/525 (6.10 %) |
| LRP1B | 2q22.1-q22.2 | 93/509 (18.27 %) | 49/525 (9.33 %) | 40/525 (7.62 %) |
