## Supplementary material for "Identification of Potential Oral Cancer-Specific Biomarker in Oral Potentially Malignant Disorders for Early Detection of Malignancy: A Systematic Review": https://acrobat.adobe.com/id/urn:aaid:sc:AP:073fb316-ae45-4717-8744-1c3f3d48940c

### Notch1

A B

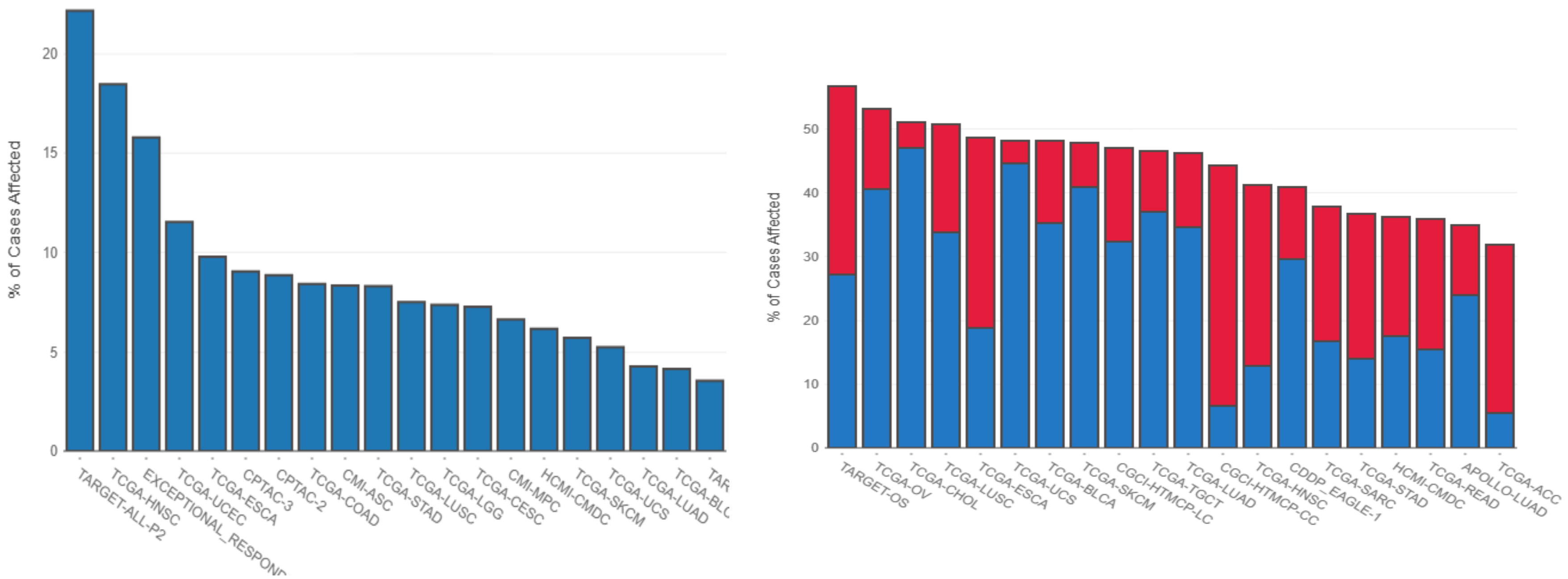


A. 812 cases affected by 864 mutations across 41 projects
B. 4,621 cases affected by 4,051 CNV events across 48 projects

# TP53

**4,964** CASES AFFECTED BY **1,355** MUTATIONS ACROSS **48** PROJECTS

**7,345** CASES AFFECTED BY **5,255** CNV EVENTS ACROSS **47** PROJECTS

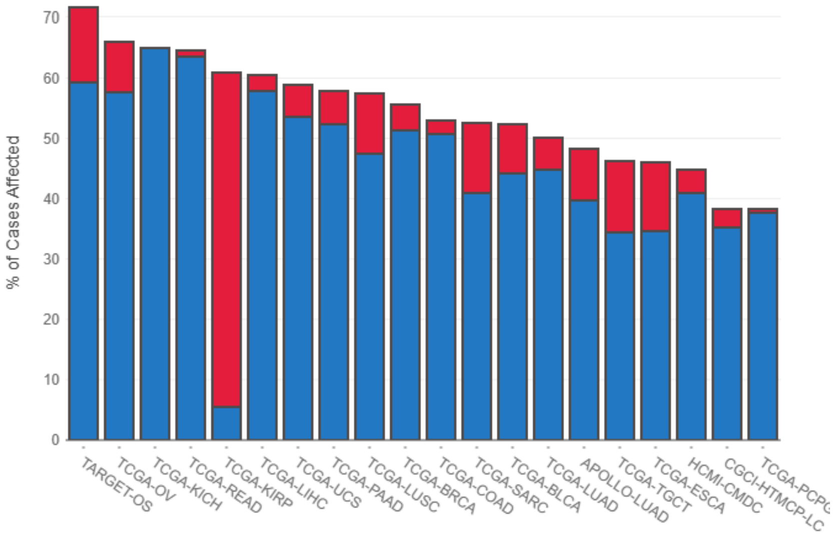


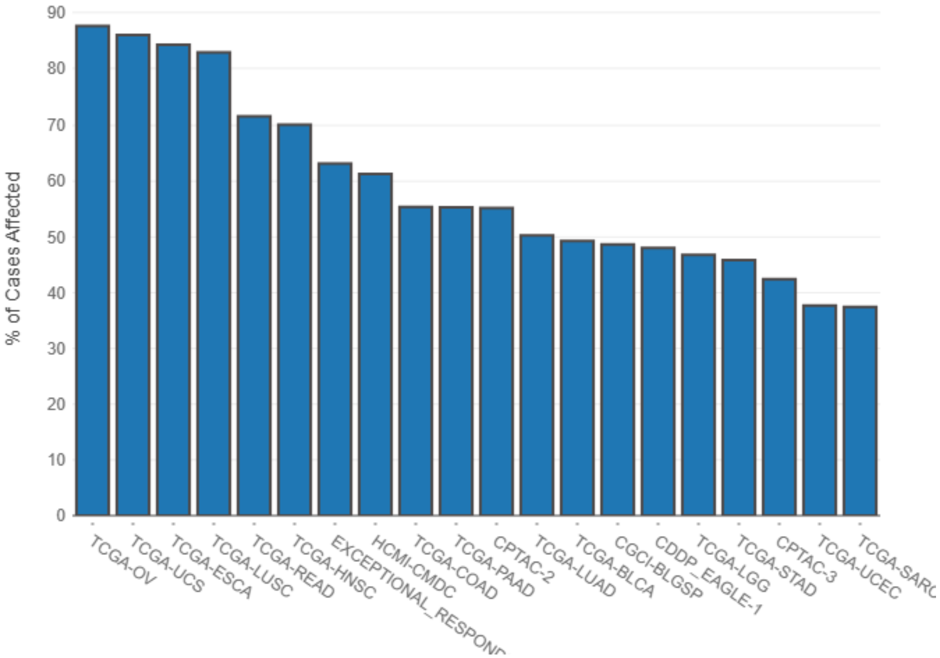

### PIK3CA

**1,685** CASES AFFECTED BY **411** MUTATIONS ACROSS **42** PROJECTS
**5,428** CASES AFFECTED BY **4,398** CNV EVENTS ACROSS **46** PROJECTS

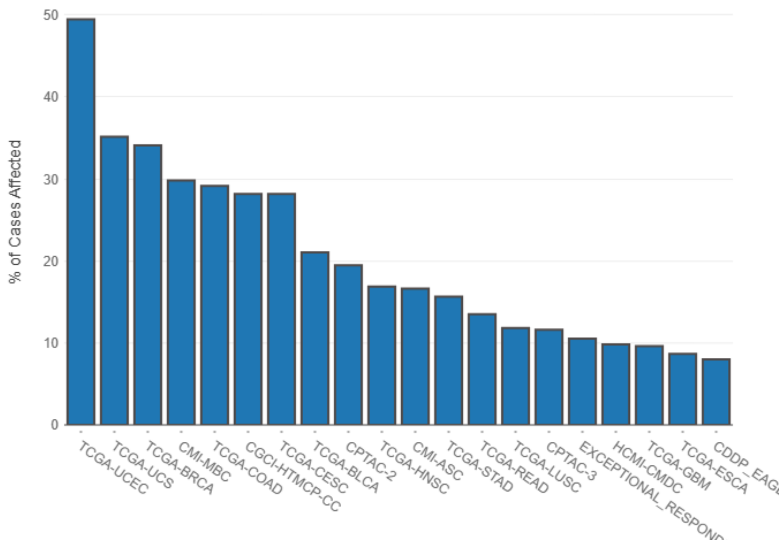


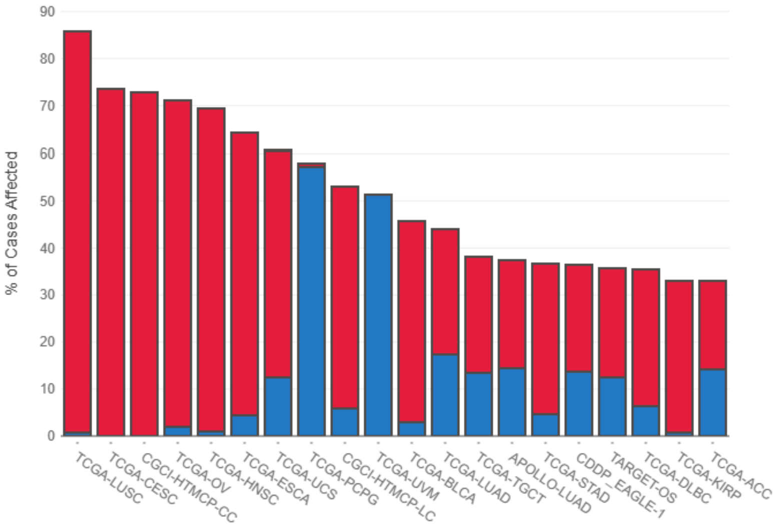

### FAT1

**1,069** CASES AFFECTED BY **1,397** MUTATIONS ACROSS **48** PROJECTS

**5,017** CASES AFFECTED BY **4,378** CNV EVENTS ACROSS **47** PROJECTS

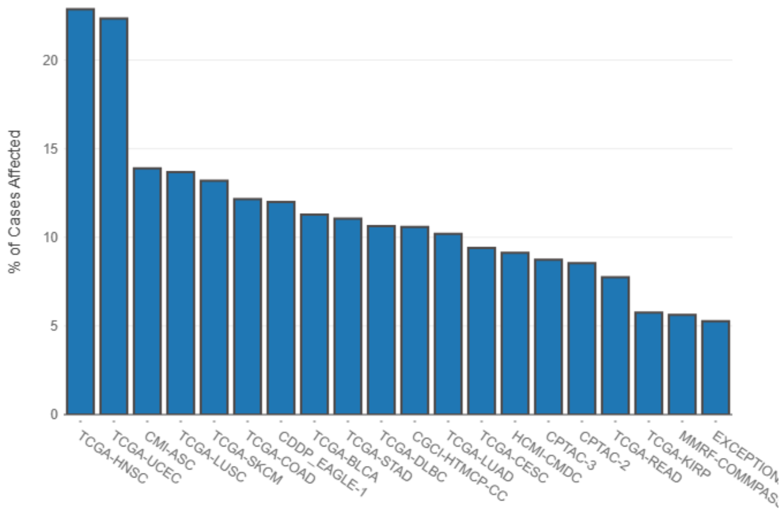


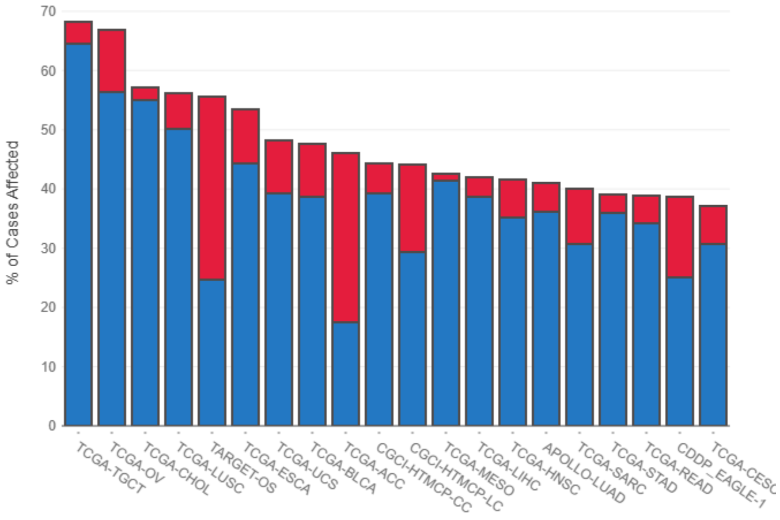

### TTN

**4,671** CASES AFFECTED BY **13,267** MUTATIONS ACROSS **50** PROJECTS
**5,929** CASES AFFECTED BY **2,805** CNV EVENTS ACROSS **46** PROJECTS

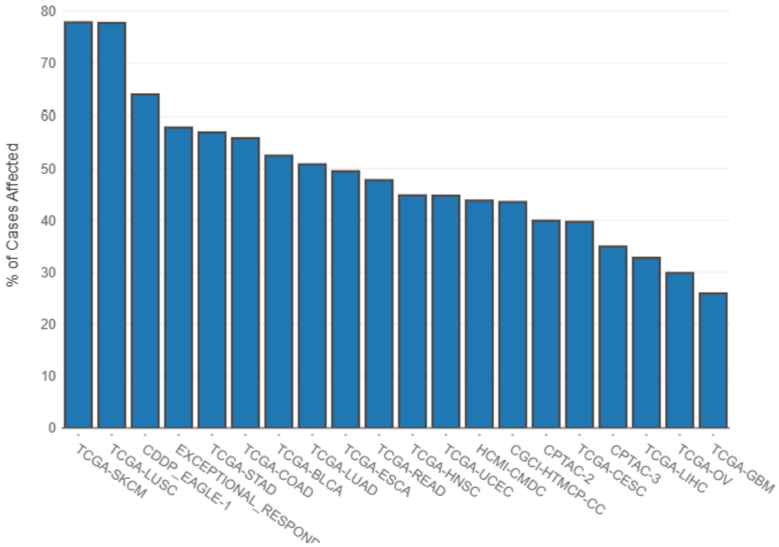


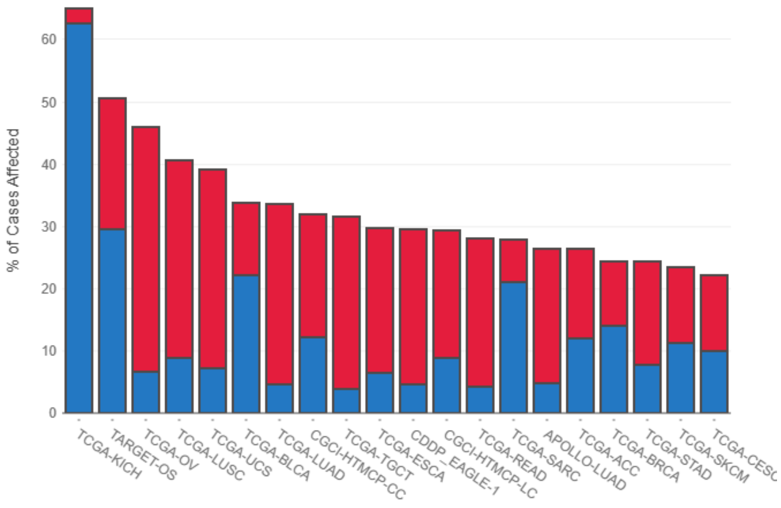

### CDNK2A

- **578** CASES AFFECTED BY **317** MUTATIONS ACROSS **34** PROJECTS

- **6,443** CASES AFFECTED BY **6,191** CNV EVENTS ACROSS **46** PROJECTS

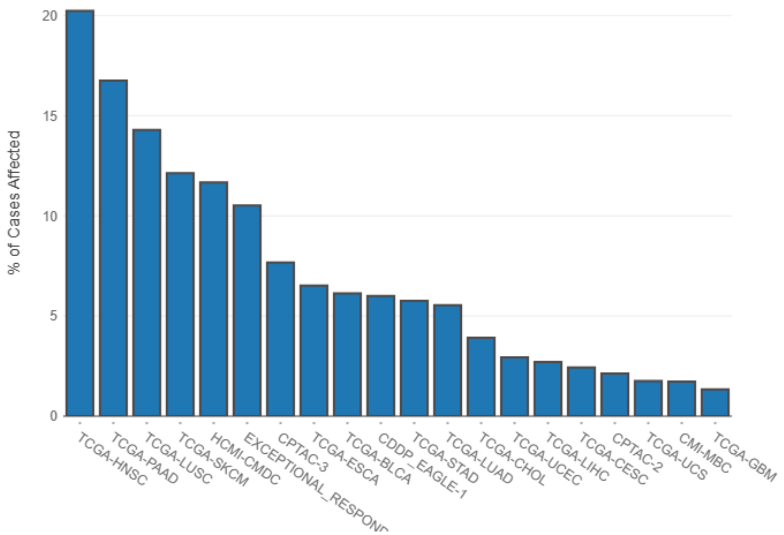


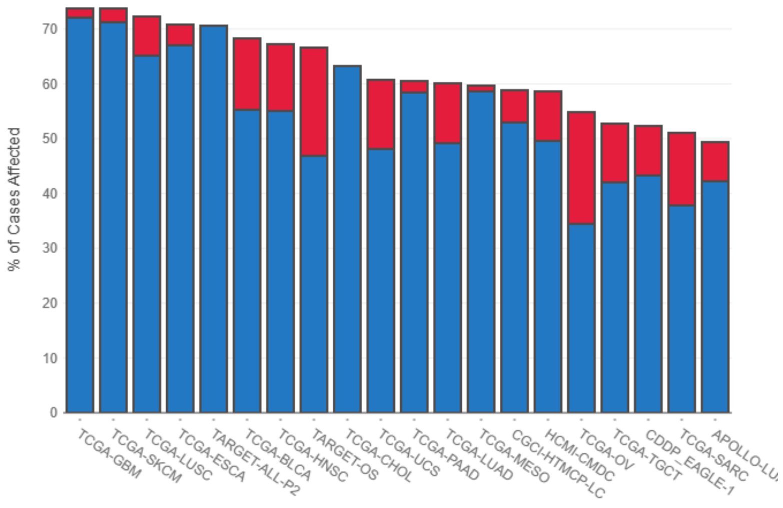

### CASP8

- **320** CASES AFFECTED BY **266** MUTATIONS ACROSS **35** PROJECTS
- **3,114** CASES AFFECTED BY **2,874** CNV EVENTS ACROSS **46** PROJECTS

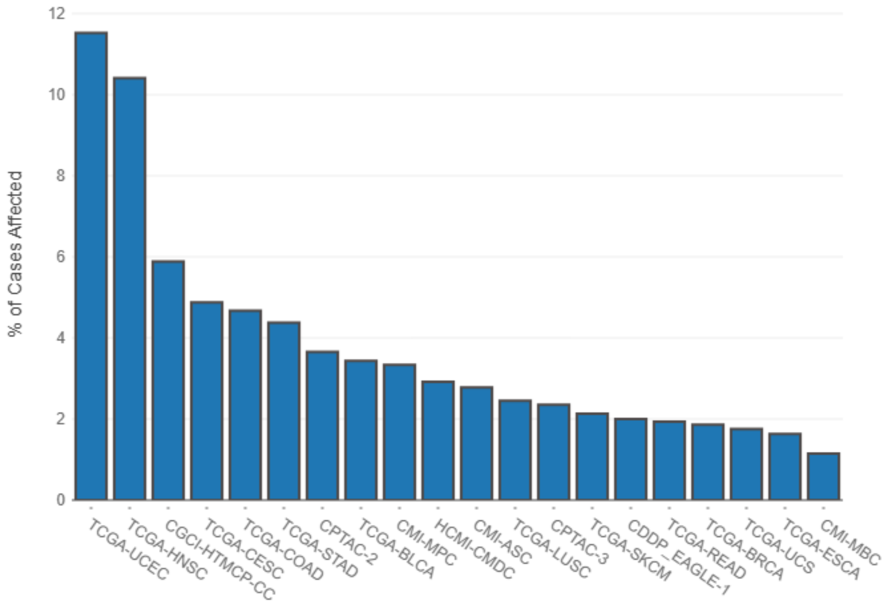


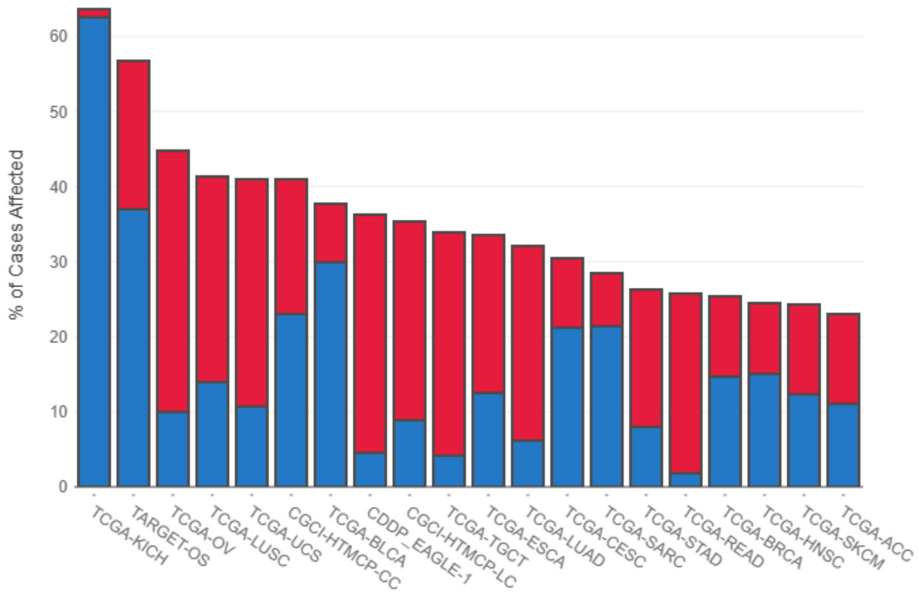

### LRP1B

- **1,784** CASES AFFECTED BY **2,697** MUTATIONS ACROSS **48** PROJECTS
- **3,860** CASES AFFECTED BY **2,685** CNV EVENTS ACROSS **47** PROJECTS

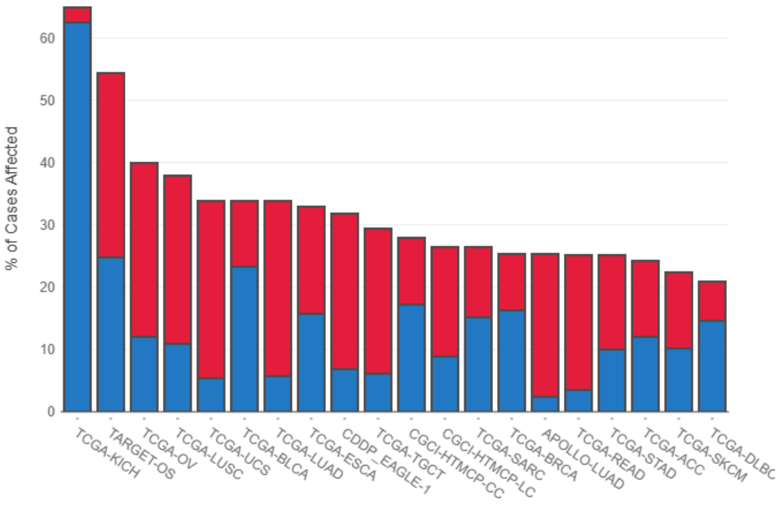


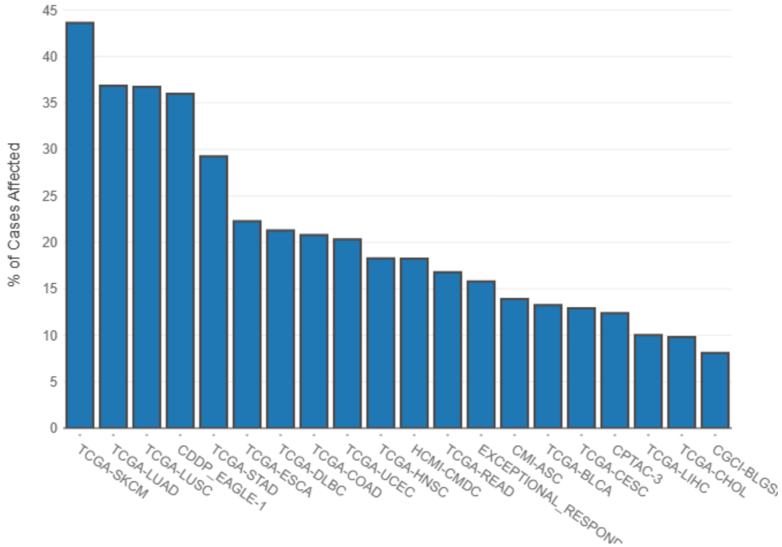
